# Water-mediated interactions between glycans are weakly repulsive and unexpectedly long-ranged

**DOI:** 10.1101/2025.02.16.638483

**Authors:** Sucheol Shin, Mauro L. Mugnai, D. Thirumalai

## Abstract

Glycans on the cell surface play an essential role in mediating cell–cell interactions as well as immune response. Despite their importance, interactions between them have not been fully characterized. Here, we reveal, using all-atom molecular dynamics simulations and free energy calculations, that water-mediated interactions between a pair of N-glycans without net charge are weakly repulsive with a range that exceeds their sizes. Unexpectedly, the effective glycan–glycan interactions decay logarithmically as the separation between them increases. Strikingly, this finding coincides exactly with the predicted interaction, which is entropic in origin, between two star polymers, consisting of long flexible polymers grafted onto colloidal particles. The weak repulsive interaction, which extends beyond the size of a glycan, is sensitive to the relative orientation of the glycans. The effective long-range repulsive interaction vanishes if the charges on water are turned off, thus establishing that electrostatic interactions, arising in part due to the persistent hydrogen bonds between water and the glycans, are responsible for the inter-glycan repulsion.

## Introduction

Naturally occurring glycans, or polysaccharides, which consist of sugar molecules joined by glycosidic bonds, are widespread biological constituents. The carbohydrate-based polymers are prevalent on the outermost surfaces of cellular or secreted macromolecules, in a variety of forms including glycoproteins where oligosaccharides are covalently linked to proteins.^1,2^ The oligomeric glycans (such as N-glycans, which are the focus in this study) play an essential role in mediating cell–cell interactions^1,3^ as well as in recognizing antigens and triggering the cascading immune responses.^4^ Especially in the context of immunology, the binding affinity of antigens to antibodies or cellular receptors could be altered significantly by modifying glycans near the binding sites.^5,6^ Although it has been long known that pathogens utilize glycans to evade the host’s immune system,^7,8^ the recent world-wide occurrence of COVID-19 brought this phenomenon into prominence.^9–12^ Because such molecular processes (binding, for example) in immune responses involve the interactions between a different variety of glycans and proteins, a systemic characterization of these interactions is needed to advance our understanding of the biological functions of glycans (such as apoptosis and cell proliferation^2^), which can be utilized to develop vaccine/antibody therapeutics^2^ and glycan-targeting drugs.^13^ Therefore, knowledge of the glycan-associated interactions is critical in the development of novel therapeutic methods through the glycan engineering of antibodies^14^ and cell surfaces.^15^

It is not straightforward to understand the interactions of glycans since their effects differ depending on the chemical context. On the one hand, glycans on a given glycoprotein could favorably interact with other proteins through non-covalent interactions, which stabilize the glycan–protein binding.^16–19^ There is a class of proteins that recognize and bind specific glycans, and the interactions between such glycan-binding proteins (GBPs) and the cognate glycans have been studied extensively. ^1,20–22^ On the other hand, the interactions between glycans could also affect the binding affinity.^23,24^ For instance, binding of an immunoglobulin G (IgG) based antibody to the cell-membrane bound receptor is strengthened by the specific glycan–glycan contacts between the antibody and the receptor.^25^ In contrast to the notion of energetic stabilization, the entropic contribution of glycans to the IgG antibody–receptor binding has also been suggested, which implies effective repulsion between glycans. ^26^

The knowledge of protein–carbohydrate interactions has increased greatly thanks to advances in the experimental techniques in the last few decades. The representative approaches include glycan microarrays^20,27,28^ and mass-spectrometry-based glycoproteomics, ^29,30^ which enable high-throughput analyses of glycan–protein binding with high sensitivity and resolution. Although these approaches are useful in identifying the binding between a glycan and a GBP, quantitative measurements of the binding constants require alternative methods such as isothermal titration calorimetry,^31^ surface plasmon resonance,^32^ and bio-layer interferometry.^33^ There have also been various experimental efforts to probe carbohydrate–carbohydrate interactions using atomic force microscopy,^34,35^ NMR spectroscopy,^36^ and hydrophilic interaction liquid chromatography.^37^ In addition, in-silico methods such as quantum mechanical calculations^18,38^ and molecular docking^39,40^ have been employed to gain microscopic insights that are not accessible solely from experiments. Most recently, molecular dynamics simulations using classical force fields^41^ have become a useful tool to predict the conformational fluctuations and atomically detailed interactions involved in a protein–glycoprotein complex, for instance, one between a virus protein and a host receptor.^10,12,42–46^

In a recent study,^47^ we investigated the consequences of entropic effect of glycans on the binding between SARS-CoV-2, the pathogenic virus of COVID-19, and angiotensin-conversion enzyme 2 (ACE2) on human cell membrane. Specifically, we calculated the binding constant for the viral spike (S) protein-ACE2 complex formation by estimating the free energy change from the number of configurations that N-glycans bonded to ACE2 adopt. Our calculations yielded nearly quantitative agreement with experiments for the change in the binding affinity upon specific glycosylation in ACE2. The assumption that S-ACE2 binding is driven by entropic forces was justified by calculating the free energy of interactions for a pair of N-glycans in water using all-atom molecular dynamics (MD) simulations.

The effective interactions between the glycans were repulsive even if polar groups in the glycans could favorably form hydrogen bonds (see Fig. S1 in the Supporting Information (SI)). Water molecules surrounding the glycans are expected to drive the repulsive interactions, as the solvent not only influences the conformational fluctuations of carbohydrates^48,49^ but also contributes to glycan–protein interactions.^50,51^ However, microscopic properties of the hydration shell of glycans and how they influence glycan interactions have not been fully explored.

In this study, we investigate the interactions between N-glycans under various conditions, which have zero net charges (i.e., neither acidic nor basic), using extensive MD simulations in explicit water and umbrella sampling calculations. ^52^ Strikingly, we find that the repulsive interactions between N-glycans, which are often branched (number of arms, *f*, does not exceed 4 in the examples considered here), are weak and long-ranged. The weak and longranged pair interactions are characterized by the orientationally averaged free energy, *G*(*r*), which varies as ln(1*/r*) for the pair distance *r* far beyond the size of a glycan (note that the logarithm changes slower than power laws, 1*/r*^*n*^). Remarkably, the MD results are in near quantitative agreement with the Witten-Pincus scaling theory for interactions between star polymers,^53^ which are models for soft colloids.^54–56^ It is surprising that despite the vast differences between glycans (small *f* and stable only in aqueous solutions) and star polymers (large *f* and likely solubilized in organic solvents) the theory for the interactions between star polymers quantitatively accounts for our findings. We also show that the glycan pair interactions are strongly repulsive if the glycans are facing each other in the opposite direction. In contrast, the interactions are weak and vary as ln(1*/r*) if the glycans are parallel to each other and oriented in the same direction. By quantifying water’s hydrogen bond structure and kinetics in the hydration shell of glycans, we highlight the role of water in mediating the inter-glycan repulsion. We establish that the effective glycan pair interactions are driven by electrostatic interactions involving water molecules as well as the glycans. In particular, in the absence of atomic charges on either water or glycans the weak logarithmic interaction between glycans vanishes. Our results rationalize the experimental observation that binding affinities of glycosylated and deglycosylated ACE2 to the SARS-CoV-2 virus are essentially identical.

## Results

### Weak and long-ranged repulsion between glycans

We simulated a pair of N-glycans by placing the system in a cubic box which also contains water and potassium chlorides at physiological concentration (Fig. 1A; see Methods for the simulation details). We considered four glycans with different sizes, GlcNAc*β*1-2Man*α*1- 6(Man*α*1-3)Man*β*1-4GlcNAc*β*1-4GlcNAc*β*1-OH, Gal*β*1-4GlcNAc*β*1-2Man*α*1-6(Man*α*1-3)Man*β*1- 4GlcNAc*β*1-4GlcNAc*β*1-OH, Gal*β*1-4GlcNAc*β*1-2Man*α*1-6(Gal*β*1-4GlcNAc*β*1-2Man*α*1-3)Man*β*1- 4GlcNAc*β*1-4GlcNAc*β*1-OH, and Gal*β*1-4GlcNAc*β*1-6(Gal*β*1-4GlcNAc*β*1-2)Man*α*1-6(Gal*β*1- 4GlcNAc*β*1-2(Gal*β*1-4GlcNAc*β*1-4)Man*α*1-3)Man*β*1-4GlcNAc*β*1-4GlcNAc*β*1-OH, which are commonly denoted as A1, A1G1, A2G2, and A4G4, respectively. These glycans have similar compositions but vary in the number of monosaccharide residues, which results in modest differences in the radii of gyration, *R*_*g*_ (Fig. 1C). From the simulations of a given glycan pair, we computed the free energy profile, *G*(*r*), by probing the pair distance *r* that is measured between the third sugar residues of the glycans (the residue numbering starts from the right in the above IUPAC names). These mannose (Man) residues are the common branching points of the N-glycans and thus regarded as the reference in comparing different glycans’ interactions.

**Figure 1:**
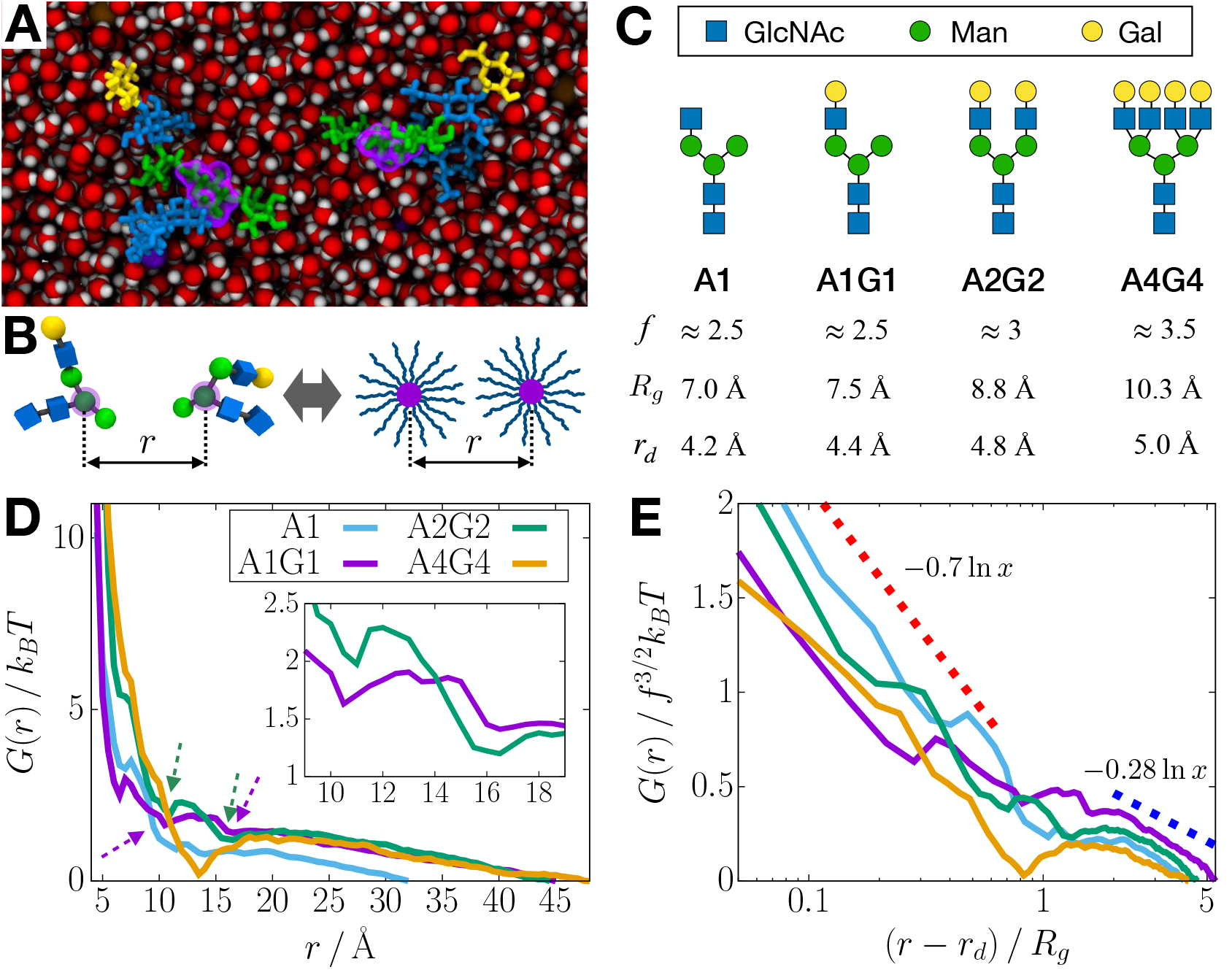
Effective interactions between charge-neutral N-glycans are weakly repulsive and long-ranged. (A) Simulation snapshot of a pair of hydrated glycans (A1G1), visualized using VMD.^57^ The glycans are represented as sticks, where the monosaccharide residues, N-acetylglucosamine (GlcNAc), mannose (Man), and galactose (Gal), are shown in blue, green, and yellow, respectively. The branch-point residues are highlighted in purple shades. Water molecules and KCl salts are in red/white and purple/brown spheres, respectively. (B) A pair of N-glycans (left) viewed as a pair of star polymers (right). The glycans are visualized using the 3D Symbol Nomenclature for Glycans (SNFG),^58,59^ where each symbol represents a monosaccharide residue. The glycan pair distance *r* is measured between the centers of mass of the branch-point residues that correspond to the core particles in star polymers. (C) Glycans with different sizes, shown in the SNFG^58^ as given on the top. For each glycan, the effective number of polymer branches, *f*, the radius of gyration, *R*_*g*_, and the lower bound of pair distance, *r*_*d*_, are shown at the bottom. (D) Free energy profile, *G*(*r*), for a pair of glycans listed in panel B, shown in units of *k*_*B*_*T*. The dashed arrows are shallow dips in *G*(*r*) for the A1G1 and A2G2 pairs, which are magnified in the inset. (E) Free energy scaled by *f* ^3*/*2^ as a function of (*r* − *r*_*d*_)*/R*_*g*_, where the *x*-axis is in a log scale. The red and blue dotted lines show the log trends with different prefactors for (*r* − *r*_*d*_)*/R*_*g*_ ≲ 0.7 and (*r* − *r*_*d*_)*/R*_*g*_ ≳ 2, respectively. Note that the prefactor 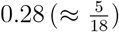 is predicted for star polymers in good solvent.

The free energy calculations showed that the interactions between a pair of glycans (A1- A1, A1G1-A1G1, A2G2-A2G2, A4G4-A4G4) are weakly repulsive, varying slowly when the inter-glycan distance (*r*) increases beyond their sizes (*R*_*g*_). In Fig. 1D, the free energy profiles, *G*(*r*)’s, indicative of weak repulsion at *r >* 2*R*_*g*_, do not change significantly until *r* ≲ *R*_*g*_. The finding that the glycans repel is consistent with the observation that oligosaccharides and some polysaccharides dissolve in water.^60^ The slowly varying nature of *G*(*r*), which is water-mediated, can be described theoretically using a scaling theory from polymer physics introduced in an entirely different context (see the next section). Note that the A1G1, A2G2, and A4G4 pairs, whose longest branches are the same, exhibit nearly identical long-ranged weak repulsion in *G*(*r*) at 20 Å ≲ *r* ≲ 45 Å. For a pair of A1 glycans, which have somewhat shorter arms, the range and magnitude of the weak repulsion is slightly smaller. This result implies that the long-range weak repulsion is related to contact between the sugar branches in a pair of glycans. Since the simulation box size for a given glycan pair is sufficiently large (see Methods for the details), the long-range weak repulsion is not a consequence of the finite size of the simulation box. Indeed, the repulsion extends even beyond the distance that is larger than twice the branch length of a glycan (for A1G1, the maximum distance from the third residue to the farthest atom in the branch is about 18 Å). Therefore, the long-range repulsion cannot arise only from the van der Waals contact between the glycan branches but is also induced by other factors such as the solvent effect and electrostatics.

At smaller values of *r*, the behavior of *G*(*r*) is best described by two distinct regimes. On the one hand, for *r* ≲ *R*_*g*_, *G*(*r*) increases steeply as *r* decreases (*i*.*e*., strong repulsion). On the other hand, for *R*_*g*_ *< r <* 20 Å, *G*(*r*) does not show significant repulsion. Notably, *G*(*r*) for the A4G4 pairs has a stabilization well with a modest depth of ~ 1*k*_*B*_*T*. The free energy profiles for A1G1 and A2G2 pairs also have shallow dips at *r* ≈ 11 Å and 16 Å (dashed arrows, Fig. 1D). Since the errors in *G*(*r*) (see Methods for the details of error estimation) are less than ~ 0.1*k*_*B*_*T*, the dips having the depth of ~ 0.2*k*_*B*_*T* are not noise. This observation suggests that the stabilization is enhanced (albeit modestly) upon an increase of glycan size due to the possibility of forming direct contact between polar groups in a glycan pair. The modest stabilization for a pair of larger glycans also aligns with the insolubility of polysaccharides in which van der Waals interactions play an important role.^61–63^ Importantly, the interactions between glycans compete against those between a glycan and water, which effectively results in the repulsive interactions. More direct evidence of the role of water in mediating interactions between glycans will be provided in a later section (see Fig. 3).

### Logarithmic pair potential—an unexpected analogy with star polymers

We found in Fig. 1D that for *r >* 2*R*_*g*_, the free energy profile *G*(*r*) is fit by the logarithm, using *G*(*r*) ~ ln(1*/r*), with *R*^2^ = 0.99. It can readily be shown that the short-range part of *G*(*r*) for *r* ≲ *R*_*g*_ is also logarithmic with a higher slope (Fig. 1E). The striking finding that *G*(*r*) ~ ln(1*/r*) is best understood using a scaling theory that was developed to predict the interaction between star polymers,^53^ which was subsequently validated using experiments and simulations.^55,64,65^ A star polymer consists of *f* flexible arms (polymers in good solvent) that are tethered to a small core, and may be thought of as a model for a soft colloidal particle.^54,55^ According to the Witten-Pincus scaling theory^53^ and the subsequent simulations^65^ by Löwen and coworkers, the interaction between a pair of two star polymers is given by,

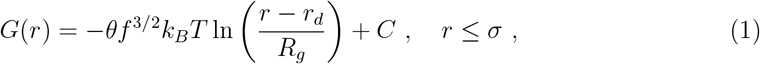

where *k*_*B*_ is the Boltzmann constant, *T* is the temperature, and *C* is an additive constant. The prefactor based on scaling arguments gives 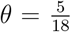.^53^ The size, *r*_*d*_, of the finite core of a star polymer may be defined as the distance at which the computed *G*(*r*) is much greater than *k*_*B*_*T*.^65^ In Eq. (1), *σ* is the corona size corresponding to the maximum range where the scaling behavior holds. The previous simulation study^65^ showed that *σ* ≈ 1.3*R*_*g*_ for the self-avoiding star polymers with various *R*_*g*_ and *f*. It is worth noting, in what is to follow, that Eq. (1) arises purely from the entropic considerations for star polymers of which *f* ≫ 1 are typical (while the original theory^53^ predicted that Eq. (1) holds for *f* = 1 and 2 as well).

We tested the applicability of Eq. (1) to a pair of N-glycans listed in Fig. 1C. The N-glycans share the similar backbone structure in which the third sugar residue is a branching point close to the glycan’s geometric center, so they may be regarded as star polymers with *f* arms (Fig. 1B). In this context, *f* is described as the *effective* number of polymer branches (or arms) at the third residue, which corresponds to the core of a star polymer. Since the branches in each glycan do not have the same number of residues, we approximate *f* using half-integers. For instance, we set *f* ≈ 2.5 for A1 that contains two branches of two sugar residues and a branch of a single residue, whereas *f* ≈ 3 for A2G2 that is more symmetric in terms of the branch length. For A4G4, which has sub-branches, we assume that the overall effect of branches on *G*(*r*) is modestly larger than for A2G2 and thus choose *f* ≈ 3.5. Indeed, the given choice of *f* values agrees with the trend in *R*_*g*_, showing that *R*_*g*_ ∝ *f*, which is reasonable. It should be stressed that regardless of how *f* is estimated for the glycans, the values are much less than for typical star polymers. We estimated *r*_*d*_ from the steep increase of *G*(*r*) for a given pair of glycans. Note that *r*_*d*_ also scales in proportional to *R*_*g*_ such that the ratio of *r*_*d*_ to *R*_*g*_ is about 0.55.

To assess if the computed *G*(*r*) has the predicted logarithmic form given in Eq. (1), we plotted the dimensionless free energy, *G*(*r*)*/f* ^3*/*2^*k*_*B*_*T*, against the reduced pair distance, (*r* − *r*_*d*_)*/R*_*g*_, with *x*-axis in a log scale (Fig. 1E). For *r* − *r*_*d*_ ≲ 0.7*R*_*g*_, the rescaled free energy profiles show qualitatively the same log trend, −0.7 ln[(*r* − *r*_*d*_)*/R*_*g*_]. The effective logarithmic dependence in Eq. (1) holds for a pair of glycans with *θ* ≈ 0.7 and *σ* ≈ *r*_*d*_ + 0.7*R*_*g*_ ≈ 1.25*R*_*g*_. The estimate of the corona size *σ* is consistent with the star polymer simulations,^65^ whereas the prefactor *θ* is ~ 2.5 times larger than expected for polymers in good solvent (*cf*. 5*/*18 ≈ 0.28). The larger prefactor implies that the repulsion between glycans at *r* ≲ 1.2*R*_*g*_ is stronger than between the long flexible polymers in good solvent. The electrostatic interactions involving both the glycans and water lead to an apparent increase in the thickness of the the glycans (see below), which explains the larger value of *θ* at short distances. For *r* − *r*_*d*_ ≳ 2*R*_*g*_, *G*(*r*) follows Eq. (1) with *θ* ≈ 0.28, and hence the glycan pairs at *r* ≳ 2.5*R*_*g*_ behave like soft colloids, demonstrating that water is a good solvent for the glycan pairs. This finding, which holds for the four different glycans in aqueous solutions, is surprising because of their small sizes and numbers of branches, *f* ‘s (max ≈ 3.5). At the intermediate distance, 1.2*R*_*g*_ ≲ *r* ≲ 2.5*R*_*g*_, the interactions between glycans exhibit modest attraction, especially for a pair of large glycans (e.g., A4G4).

### Interactions between glycans are strongly dependent on orientation

An N-glycan adopts a specific geometry in its glycosidic bond to the glycosylation site of a given protein such that its first monosaccharide residue (GlcNAc) is linked through the amide nitrogen of an asparagine residue on the protein. Hence, the interactions between N-glycans are expected to depend on how they are oriented to each other. For example, the N-glycans on ACE2, the receptor protein of the SARS-CoV-2 virus, protrude from the protein surface and their relative orientation is roughly in the parallel geometry (see Fig. S2). On the other hand, when the glycosylated spike (S) protein of the virus approaches ACE2 for binding, the S-glycans are likely to encounter the ACE2-glycans in the anti-parallel orientation. The glycan–glycan interactions corresponding to these two different geometries are distinct. Therefore, it is important to calculate the dependence of the free energy profiles on the relative orientation of the glycans.

To determine the orientation dependence, we computed the free energy profile for a glycan pair that is orientationally constrained. We considered two extreme cases, where the glycans face each other in parallel and opposite (or anti-parallel) directions (Fig. 2A), which are denoted by ↑↑ and →←, respectively, using the arrow to indicate the direction of each glycan’s sequence. We constrained the glycans in the simulations such that the vectors connecting residue 1 to residue 3 were oriented in either parallel or opposite directions. A harmonic spring potential with the varying equilibrium distance was applied on the central residues for the umbrella sampling (see Fig. 2A). More detailed description for the constraints imposed in the simulations is provided in Methods.

**Figure 2:**
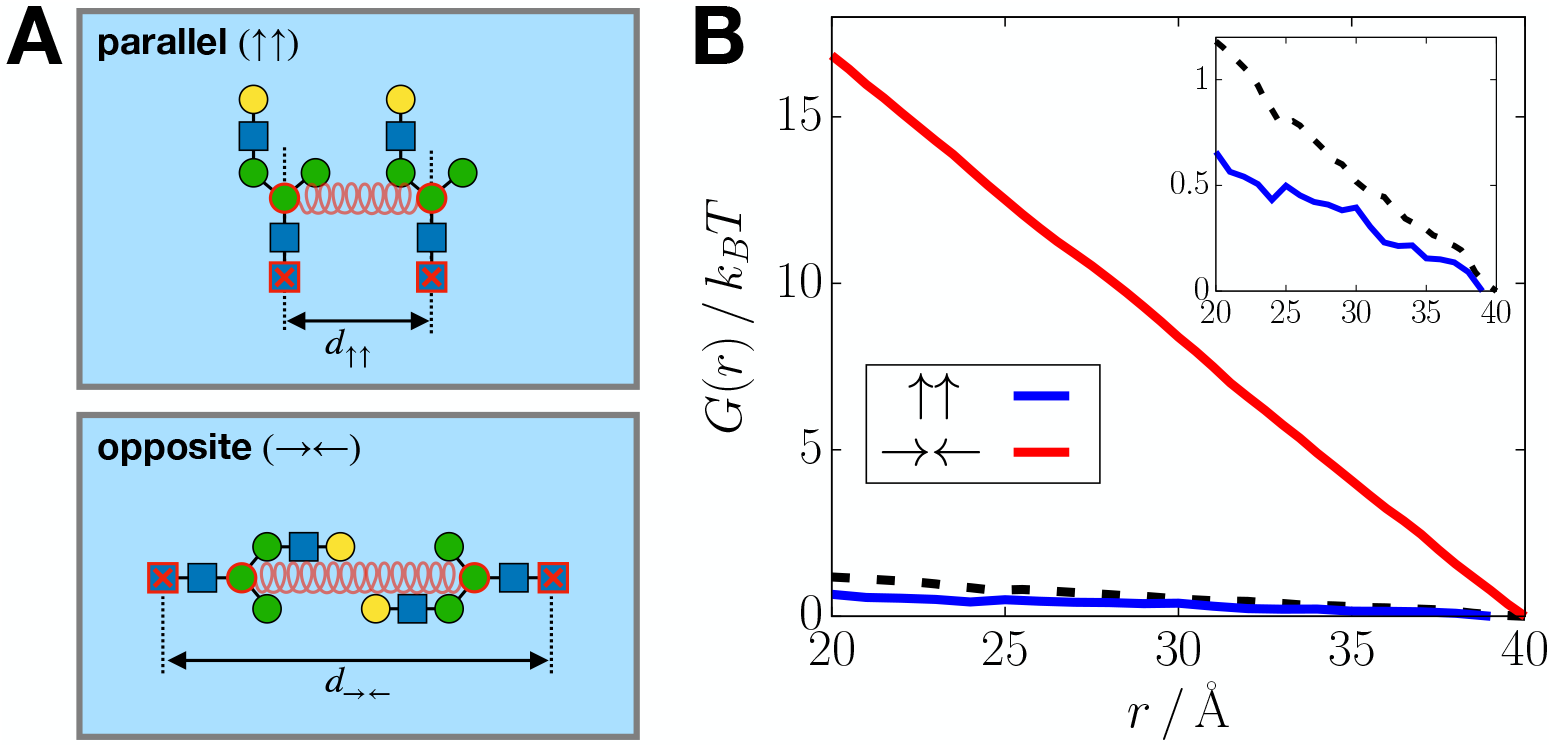
Repulsive effective interactions between glycans are enhanced when their branches are oriented to each other. (A) Schematic illustrations of the simulated systems for a pair of A1G1 in water with parallel (top) and opposite (bottom) orientations. The residues with red borders are constrained to maintain specific orientation in the simulations. The first residues (GlcNAc; blue squares with red crosses) are restrained at the points separated by a given distance (*d*_↑↑_ or *d*_→←_), and the third residues (Man; green circles) are subject to a harmonic spring potential. (B) Long-distance part of *G*(*r*) for a pair of the glycans constrained in the parallel (↑↑) and the opposite (→←) orientations. The black dashed line is *G*(*r*) obtained without the pair orientational constraint. The inset shows the enlarged view of the data for the parallel orientation in comparison with the constraint-free case.

In Fig. 2B, the resulting free energy curves at *r* ≥ 20 Å are shown for the glycan pair constrained in either parallel (↑↑) or opposite (→←) orientations. Although the interactions between the glycans are repulsive in both the orientational constraints, the extent of repulsive interactions is strikingly dependent on the orientation. When the glycan pair is oriented in the ↑↑ geometry (Fig. 2A, top), the slope of the free energy curve decreases to about a half of the case without constraint (Fig. 2B, blue). On the other hand, for the glycan pair oriented as in →← (Fig. 2A, bottom), the effective pair repulsion becomes even stronger showing the slope that is about 15 times larger than the orientationally averaged *G*(*r*) (Fig. 2B, red). These results highlight that glycans experience strong repulsion when their branches are in proximity. Hence, such glycan–glycan contacts in the opposite orientation would be minimized, for instance, in the binding process of the SARS-CoV-2 virus to ACE2. Once the complex is formed between the viral spike protein and ACE2, glycans around the binding interface are mostly in the orientation close to the anti-parallel but staggered (i.e., not colinear) geometry such that the interactions between the glycans are only weakly repulsive.

### Role of water in the inter-glycan repulsion

#### Structure of water is unaffected by glycans

The orientation-averaged interactions (Fig. 1) show that the hydrated glycans behave like star polymers in good solvent. To understand the role of water in mediating the repulsive interactions between glycans, we characterized the structural and dynamical properties of water’s hydrogen bonding in the hydration shell of the glycan, A1G1. First, using the simulation of a single hydrated A1G1, we calculated the tetrahedral order parameter, *q*_tet_, which represents the structure of hydrogen bond (HB) network in water^66^ (see Methods). The distribution of *q*_tet_ in Fig. 3A does not show a significant difference between water molecules close to the glycan and those in the bulk liquid phase, reflecting that the HB network of water around the glycan is nearly identical to that in the bulk water. If we consider *q*_tet_ for water molecules that are only adjacent to the glycan’s oxygen or nitrogen atoms, then the distribution shifts toward the right such that the mean value increases modestly from 0.55 to 0.58. Hence, water molecules in the glycan’s first hydration shell are deemed to have approximately the same extent of tetrahedral HB network as those in the bulk, which would make a given water molecule favor being at the glycan surface to the same extent as in the bulk liquid. Therefore, direct contact between glycans would be disfavored by water molecules in the solvation shell. In other words, water is a good solvent for these glycans. Indeed, the distribution of *q*_tet_ for water molecules in between a pair of A1G1 is shifted toward the left from that around a single A1G1 (Fig. 3A). Although the difference is not large in terms of the mean (0.53 vs. 0.55), this result also provides a reasonable basis for how water makes glycans repel each other.

**Figure 3:**
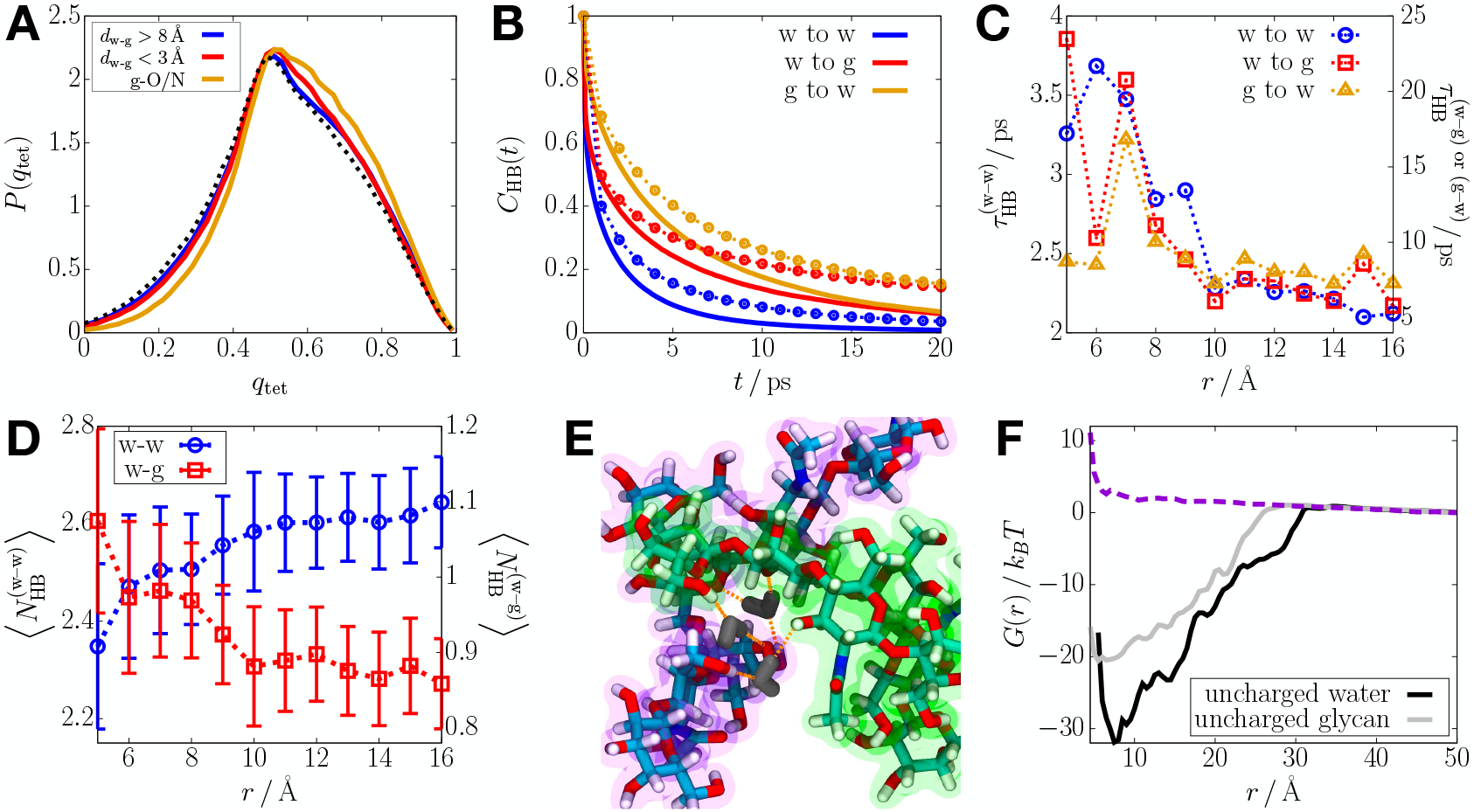
Water mediates the effective repulsive interactions between glycans. (A) Distribution of tetrahedral order parameter for water’s HB network in the bulk liquid phase (blue), near A1G1 (red), and particularly near the oxygen or nitrogen atoms of the glycan (orange), where *d*_w-g_ is the distance between a given water molecule and the glycan. The dotted line is for water molecules in between the A1G1 pair constrained to *r* = 10 Å. (B) Time correlation function of a hydrogen bond (HB) formed between water molecules (blue), donated from water to the glycan (red), and donated from the glycan to water (orange). The solid lines show the correlation functions for the HBs in the hydration shell of a single glycan, whereas the dotted lines with circles are from the HBs of water in between the glycan pair constrained to *r* = 7 Å. (C) HB lifetime for water in between the glycan pair as a function of *r*. The left side of *y* axes reads the values for the blue data points, whereas the right side reads the red and orange. (D) Average number of HBs per water molecule in between the glycan pair as a function of *r*. The left and right sides of *y* axes read the blue and red data points, respectively. (E) Simulation snapshot illustrating water molecules (dark-gray) forming hydrogen bonds (orange dotted lines) with the glycan pair (green and purple shades) at *r* = 7 Å. (F) *G*(*r*) for an A1G1 pair in uncharged water (black) in comparison with that in normally charged water (purple dashed). The gray curve shows *G*(*r*) for the uncharged glycans in normal water.

#### Hydrogen bond kinetics

Next, we analyzed the kinetics of hydrogen bond relaxation around a single A1G1 by computing the time correlation function, *C*_HB_(*t*) (see Methods). We found that a HB formed between water molecules near the glycan decays faster than one between water and the glycan (Fig. 3B). The relaxation times for the former (HB between water) and the latter (HB between water and glycan) are 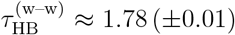 ps (*cf*. ≈ 1.1 ps in the bulk liquid) versus 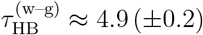 ps (for a HB from water to the glycan) or *τ* ^(g–w)^ ≈ (±0.4) ps (for a HB from the glycan to water). The longer lifetime of a HB involving the glycan implies the tendency for a water molecule to stick around the glycan. 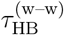 increases modestly to 1.97 (±0.02) ps when the water molecule is in the vicinity of the glycan’s oxygen or nitrogen atom that can donate (accept) a HB to (from) water (Fig. S3A). The direct hydrogen bonding interactions between water and glycan are thus likely to stabilize HBs between water molecules in the hydration shell, which is consistent with the modest increase of *q*_tet_ for water molecules near the glycan’s oxygen or nitrogen atoms (Fig. 3A).

For the hydration shell in between a pair of A1G1, the HB kinetics is slower than for a single A1G1. Especially when a pair of glycans are so close to each other (*r* ≲ *R*_*g*_), a HB formed between water and the glycan is more stable as reflected in the four-fold increase in the HB lifetime (Figs. 3B and 3C). In addition, 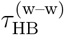 shows a twofold increase at the same time. *C*_HB_(*t*) for the glycan pair at small separation shows the long-time decay that is much slower than for a single glycan (*t*^−0.7^ vs. *t*^−2.5^ for *t >* 10 ps; see Fig. S3B), which reflects fundamental difference in the hydrogen bond environment. Interestingly, water molecules tend to form hydrogen bonds more with glycans than with other water molecules upon decreasing *r* (Fig. 3D). Note that the number of hydrogen bonds for the sandwiched water with glycans fluctuates around 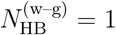, suggestive of two hydrogen bonds formed between water and the glycan pair with significant probability (Fig. 3E). Hence, it would be more difficult to remove water molecules that are hydrogen-bonded to the glycans at short distances. Such discrete water molecules that are strongly bound to the glycans, effectively making the glycan arms thicker, contribute to the repulsion between glycans at small *r*, observed in Fig. 1E. Interestingly, the results reported herein are comparable with those for a highly concentrated solution of simple monosaccharides (e.g., fructose)^67,68^ in which the sugar molecules form a percolated cluster through water.^69,70^

### Glycan–water versus glycan–glycan interactions

We also analyzed the structural and dynamical properties of water’s hydrogen bonds in the hydration shell of A4G4 that exhibits a shallow attraction well in the pair free energy profile (Fig. 1D). We found that the distribution of *q*_tet_ for a single A4G4 is indistinguishable from that for a single A1G1 (Fig. S4A). For the water molecules that are in between a pair of A4G4, the *q*_tet_ distribution shows a leftward shift similar to that for the A1G1 pair. The changes in HB lifetimes and number of HBs, upon decreasing *r*, are qualitatively comparable between A1G1 and A4G4 (Figs. S4B and S4C). It is notable that the HB lifetimes for the A4G4 pair are systematically longer than for the A1G1 pair, which may be attributed to the increased polar interactions involving the oxygen or nitrogen atoms of A4G4 due to its larger size. Nevertheless, these properties of water do not account for the modest attraction between the A4G4 pair, which arises at 12 Å ≲ *r* ≲ 18 Å in *G*(*r*) (Fig. 1D). Rather, we found that the modest stabilization for the A4G4 pair is associated with the significant increase in the number of HBs directly formed between the glycans, 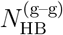, compared to that for A1G1. The average number of inter-glycan HBs increases by ~ 11.5 times at *r* = 14 Å where *G*(*r*) for the A4G4 pair reaches the minimum (Fig. S4D). Remarkably, the number of HBs decreases sharply at *r* = 12 Å below which the effective glycan interactions are largely repulsive. Although more HBs are likely to form upon decreasing *r*, relatively much larger increase in 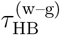 would lead to the effective repulsion for A4G4 at a small pair distance as a result of the competition between glycan–glycan and glycan–water interactions.

#### Importance of electrostatic interactions

To directly assess the contributions of glycan– water and glycan–glycan interactions to the free energy profile, we repeated the free energy calculations using systems with modified electrostatics in water and glycans (see Methods for the simulation details). First, we computed *G*(*r*) for the A1G1 pair in the artificial solvent of *uncharged* water. The calculated free energy profile shows that the effective pair interactions are favorable over a wide range of pair distances (6 Å ≲ *r* ≲ 30 Å) with stabilization up to ~ 32*k*_*B*_*T*, in contrast to *G*(*r*) for the glycans in the normal water which is almost monotonically increasing as *r* decreases (Fig. 3F). The large difference in *G*(*r*) highlights the central role of water in the effective repulsion between the hydrated glycans, since the bare glycans without water are attractive to each other due to the favorable electrostatic interactions (e.g., HB-like) between them. Interestingly, treating water as a dielectric continuum does not recover the effective repulsion between glycans (Fig. S5), which shows the importance of hydrogen bond network of water in driving the repulsion. On the other hand, the weakly repulsive tail of *G*(*r*) at longer distances, *r* ≳ 35 Å, is comparable to that for the hydrated glycans (Fig. 3F). The repulsive tail does not change either upon rendering the glycans uncharged in normal water (gray curve, Fig. 3F; note that the favorable *G*(*r*) reflects the attraction between non-polar hydrophobic moieties in water). These observations, however, do not rule out the electrostatic contributions of water and glycans to the repulsion at large *r*, because *G*(*r*) for the glycans in vacuum shows a larger extent of repulsion at *r* ≳ 35 Å than in water (Fig. S5). It is thus suggested that the electrostatic interactions of both water and glycans also play a delicate role in rendering glycans to have the long-range weak repulsion.

## Discussion

We used molecular dynamics simulations to show that water-mediated effective repulsive interactions between charge-neutral N-glycans are unusually long-ranged, extending beyond their sizes (not just beyond *R*_*g*_ but also farther than twice the branch length of a glycan). Surprisingly, the free energy profiles, characterizing the effective glycan–glycan interactions, vary only as *G*(*r*) ~ ln(1*/r*), a result that was established to describe entropically-driven interactions between star polymers in good solvent. The origin and context of the effective interactions between glycans in water and star polymers are entirely different. In idealized star polymers, typically dissolved in organic solvent, *f* flexible polymers are grafted with one end of each chain that is covalently attached to a sphere, referred to as a corona. The interactions between the polymers and between the coronas are weakly repulsive, varying as *G*(*r*) ~ ln(1*/r*). In contrast, N-glycans, which are polar and hydrophilic, are solvated in a highly dielectric medium. Nevertheless, the predicted logarithmic interaction between two star polymers, nominally with *f* ≫ 1, also holds for glycans with small *f*, which is indeed striking and unexpected. The weak logarithmic interaction between two star polymers is the result of a reduction in the number of configurations that the grafted polymers can adopt at distances that are short compared to size of the polymer chains.^53^ In other words, the prediction *G*(*r*) ~ ln(1*/r*) is entropic in origin. In contrast, we have shown that electrostatic interactions, especially involving water, plays an important role in the origin of the effective repulsion between glycans. The surprising finding that the theory for the interactions between star polymers is in accord with the interactions between glycans could also be relevant in a variety of oligomeric/polymeric molecules that are polar but non-ionic. For instance, water-soluble block copolymer micelles^64,71^ (which may be regarded as star polymers^56^) or hydrophilic polymer brushes^72,73^ could exhibit effective repulsion in water like the inter-glycan interactions. This concept may also be applicable to the interactions involved in a broad class of glycopolymers.^74,75^ Importantly, our work also shows that there could be multiple mechanisms for the origin of the logarithmic interactions, which implies that it is more prevalent than previously thought (as highlighted in the experiments^35^ to be discussed below).

Although the main results may be rationalized by theoretical arguments based on an analogy to star polymers^53^ as well as the empirical observation of oligosaccharides’ solubility in water,^60^ it is desirable to provide plausible validation by appealing to experiments. One possible way to test the predicted logarithmic dependence of glycan–glycan interactions would be to make use of atomic force microscopy (AFM) where the probes are tagged with oligosaccharides. Previously, experiments using AFM quantified carbohydrate–carbohydrate interactions pertinent to cell–cell adhesion. ^34,35^ These experiments mostly focused on the attractive forces in the range of ~ (200–300) pN between negatively charged carbohydrates.^34^ To further rationalize our results, we compared to the measured force between the colloidal probes decorated with charge-neutral disaccharides in the absence of Ca^2+^.^35^ In this case, the force profile is purely repulsive over a wide range of the probe distances (Fig. 4, left panel), which is consistent with our prediction based on the simulations. The range of the force magnitude also agrees with that computed from the simulated free energy (i.e., *F* (*r*) = −*dG*(*r*)*/dr*). Notably, by re-plotting the experimental force profile in a log scale (Fig. 4, right panel), we found that the repulsive force between the carbohydrate-coated probes is inversely proportional to the probe distance (i.e., *F* (*r*) ~ 1*/r*), which implies the logarithmic interaction, *G*(*r*) ~ ln(1*/r*). The range of the logarithmic repulsion in the experiment (~ 40 nm) is longer than in the simulations (~ 5 nm), which may be attributed to the thiol linker used to attach the carbohydrates to the probe as well as the long-range interactions between the lipid membrane-coated probes.^35,76^ Overall, the experimental results support our simulation-based findings to some extent. Additional experiments for a number of glycans along with the computational investigations would be able to provide further validation.

**Figure 4:**
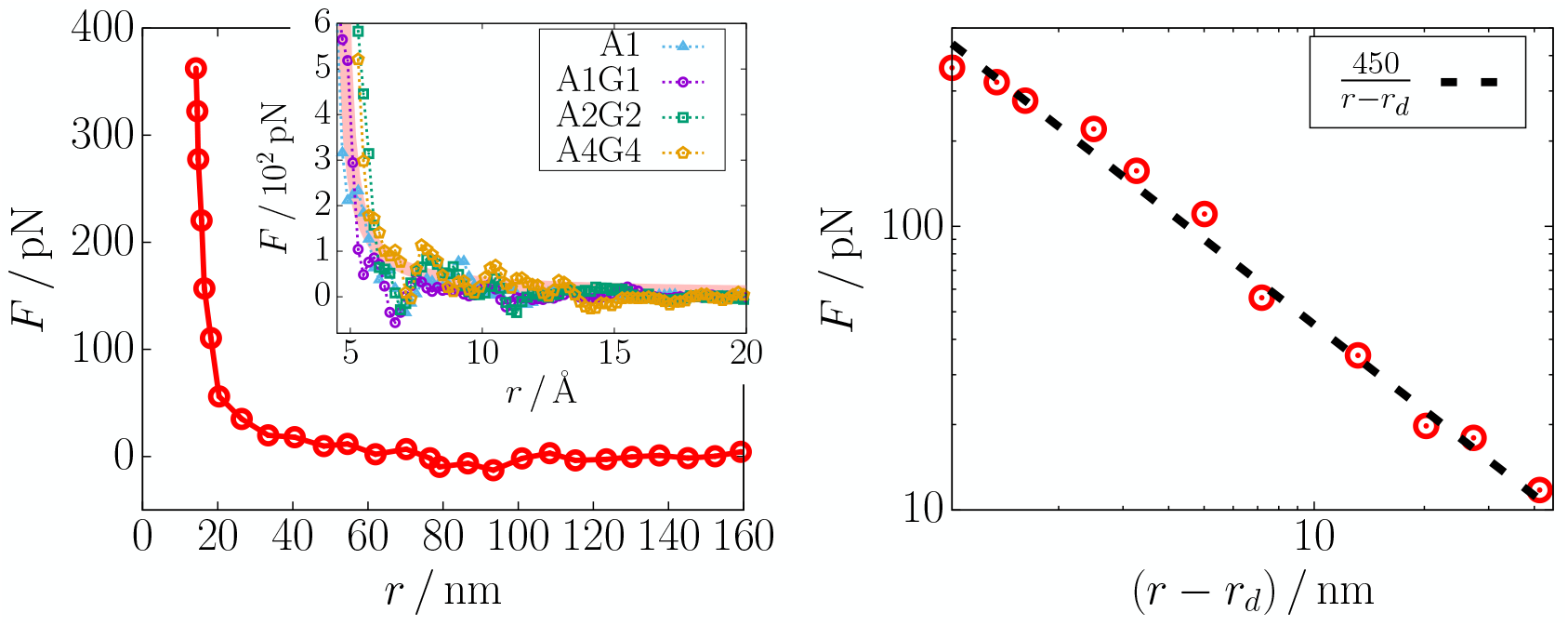
Glycan–glycan interactions measured using atomic force microscopy support the logarithmic dependence on the probe distance. (left) Force profile for a pair of charge-neutral disaccharides covalently linked to colloidal probes in the EDTA buffer without Ca^2+^,^35^ where the data points were extracted from Figure 1b in Ref. 35 using WebPlotDigitizer. ^77^ The inset shows the mean force computed from the simulated free energy profile for different glycan pairs, where the pink solid line illustrates the overall trend of *F* ~ 1*/r*. (right) Log-log plot of the experimental force profile where the distance is shifted by *r*_*d*_ = 13.2 nm. The black dashed line corresponds to *F* ~ 1*/r*. The inverse relationship between the force and the distance, *F* ~ 1*/r*, corresponds to *G*(*r*) ~ ln(1*/r*).

The major finding reported here has implications for the post-translation process of glycosylation, which involves attachment of glycans to sites on a protein. Although known for a long time, glycosylation comes into sharp focus because of its role in the recognition of human cell by the receptor binding domain (RBD) of the virus SARS-CoV-2, which engages with the peptidase domain (PD) of the cell membrane protein, referred to as angiotensin-converting enzyme 2 (ACE2).^78,79^ The PD has six asparagines that are susceptible to glycosylation. Interestingly, experiments showed that modification of the glycoform of ACE2 did not affect RBD binding.^80–83^ In a previous study,^47^ we showed that the experimental measurements could be quantitatively explained by assuming that the effect of ACE2 glycans on the RBD binding is entropic in origin. The results of the current study show that water-mediated interactions between glycans are weakly repulsive, thus justifying the assumption that ACE2 glycans exert only weak repulsion on the glycans near the RBD in the viral spike protein. This prediction could be tested by studying the temperature dependence of ACE2 binding to RBD.

The dramatic orientation dependence of glycan–glycan interactions may also have several implications. First, the parallel arrangement of glycans (Fig. 2A), likely to occur on a single glycoprotein surface, would enhance the stability of the folded protein ^1,84,85^ by a molecular crowding (entropic) effect,^86^ since the repulsive interactions in the parallel geometry are negligible. If the interactions were too strong between adjacent glycans on a given glycoprotein, they would impose a mechanical strain along the protein surface, which could adversely impact the protein’s folding. ^87–89^ In contrast, glycans oriented in the opposite geometry (Fig. 2A), which may occur in between distinct glycoproteins approaching towards each other, are subject to strong repulsion. Such unfavorable interactions would be avoided and thus contribute to the solubility of glycoproteins. ^1^ In this context, glycans on a pathogen can serve as the “shield” or “camouflage” against the immune system.^9^ For SARS-CoV-2, while almost the entire spike protein is shielded by glycans, the certain protein conformations in which RBD is not covered would enable binding with ACE2. ^12,90^ In the RBD-ACE2 complex, glycans near the binding interface would interact mostly in the anti-parallel (staggered) geometry, exerting only weak repulsion on one another. Therefore, the contribution of glycan–glycan interactions to the RBD-ACE2 binding affinity is entropic and hence not significant in magnitude. Since this argument is general, it should be applicable to other binary systems involving interactions between charge-neutral glycans.

Like many others, our study has a limited scope. A major limitation is that the findings and implications addressed above are restricted to N-glycans that are electrically neutral. For water-insoluble polysaccharides (e.g., chitin and cellulose), van der Waals (vdW) interactions, in addition to the inter-carbohydrate hydrogen bonds, significantly contribute to the stabilization of the carbohydrates.^61–63^ In contrast, N-glycans have much shorter chain lengths such that the vdW stabilization may be less important than the hydrophilic interactions between the glycans and water, which results in an effective repulsion between the glycans and renders it water-soluble. The effects of vdW interactions, albeit not fully explored here, would need to be considered more carefully when studying the interactions between glycans with much longer chain lengths such as glycosaminoglycans (GAGs). Since GAGs are negatively charged due to their acidic groups (e.g., sulfate or carboxylate), the effects of charges and ions would need to be addressed as well.^91,92^ Another weakness of our study is that it does not explicitly take into account the potential effects of glycan–peptide interactions^17,93^ on glycan–glycan interactions in the binding between a virus protein and a glycosylated receptor protein. Future investigations of such effects would further elucidate the roles of glycan-associated interactions in biological processes.

## Methods

### MD simulations for umbrella sampling

We performed all-atom MD simulations for glycans in explicit water using the CHARMM36 force field.^94^ Two N-glycans of the same kind (listed in Fig. 1B) were solvated in a cubic box in TIP3P water^95,96^ in the presence of 150 mM KCl. The glycans were subject to a biasing potential, 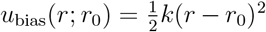, which constrains the pair distance *r* to fluctuate around *r*_0_. The harmonic spring force with *k* = 1.8 kcal*/*(mol · Å^2^) was applied between the centers of mass for the third residues of two glycans. The initial configurations of the hydrated glycans at varying *r*_0_ with the increment of Δ*r*_0_ = 1 Å were prepared. The side length of the simulation box, *L*, was chosen such that *L >* 2*r*_0_, to ensure that the effect of finite size of the simulation box is minimal. Each system was subject to 2,000 iterations of energy minimization based on the conjugate gradient method and the equilibration for 1 ns at 300 K and 1 bar using a Nose-Hoover thermostat^97^ and a Parrinello-Rahman barostat.^98^ The glycan pair distance was sampled every 0.2 ps in the subsequent production run of 50 ns (for *L <* 100 Å) or 30 ns (for *L* = 100 Å) under the same NPT condition. We constructed the free energy curve, *G*(*r*), by applying the standard Weighted Histogram Analysis Method (WHAM)^99^ to the glycan pair distances sampled from the simulations. The error associated with the free energy was estimated from 200 iterations of the WHAM calculations using the samples randomly reconstructed from the original statistics. The selected samples for each umbrella window are uncorrelated (i.e., the temporal separation between the samples is larger than the correlation time, ~ 5 ps). The calculations of *G*(*r*) and the errors were performed using the publicly available code.^100^

For the dynamic propagation of a system, the velocity Verlet algorithm was used with the integration time step of 2 fs. The geometry of water and all the bonds involving hydrogens were restrained using the SHAKE algorithm.^101^ The Particle Mesh Ewald method with periodic boundary conditions was used to compute the long-range part of electrostatic interactions.^102^ All energy minimization and simulations were performed using the molecular dynamics packages, LAMMPS^103^ and NAMD.^104^

### Umbrella sampling for the glycans with orientational constraint

As illustrated in Fig. 2A, we sampled the configurations of the glycan pair, which is subject to the biasing potential described above, with constrained relative pair orientation. For the parallel orientation, the group of atoms in the first residues (GlcNAc) were pinned, respectively, at two spatial points separated by *d*_↑↑_ = *r*_0_ where *r*_0_ is the equilibrium distance for the biasing potential imposed on the third residue. Harmonic spring potentials with *k* = 300 kcal*/*(mol · Å^2^) were used for the pinning. A cubic box with *L* = 60 Å was used for the simulation cell, and the initial configurations with various *r*_0_ were prepared by placing the parallel-oriented glycan pair along the diagonal of the simulation box.

For the opposite orientation, the first residues were restrained around the points with 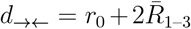, where *R*_1–3_ is the distance between the centers of mass of the first and the third residues and the bar denotes an average. Since the distribution of *R*_1–3_ has a longer tail toward the left side due to the chain’s flexibility (see Fig. S6A), we took the median for the average, which gives 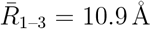. Less stiff spring potentials with *k* = 0.5 kcal*/*(mol · Å^2^) were used to allow the first residues to fluctuate around the fixed points by ≈ 1 Å. A was used for the simulation cell where rectangular box with dimensions of 40 × 40 × 90 Å^3^ the glycans were placed along *z* axis in the opposite and collinear direction for the initial configurations. All other simulation conditions were kept the same as described in the previous subsection (i.e., without orientational constraint), except that the statistics were generated from the production run for 20 ns. The distribution of the relative orientation between a glycan pair constrained as above shows a predominant peak at the parallel or opposite sense (Fig. S6B).

### Simulations with modified electrostatics

To generate the free energy curves shown in Fig. 3F, we performed umbrella sampling for the A1G1 pair with modified atomic charges in the system. For the glycans in the explicit solvent of uncharged water, the atomic charges on the water molecules were set to zero while the geometry and Lennard-Jones (LJ) interactions were unaltered. The ions were also treated as charge-neutral LJ particles such that the electrostatic interactions only occur between the glycans. The volume of the simulation box was fixed to maintain the solvent density as in normal water. The harmonic biasing potential was imposed with the same spring constant as above, *k* = 1.8 kcal*/*(mol · Å^2^). The configurations corresponding to *r*_0_ *<* 25 Å and *r*_0_ ≥ 25 Å were sampled from the simulations with *L* = 60 Å and *L* = 100 Å, respectively. For the uncharged glycans in explicit water, all the conditions were kept the same except the glycan’s atomic charges which were set to zero. For the free energy curves in Fig. S5, we performed umbrella sampling for the glycan pair without any solvent and ions. The umbrella sampling calculations were repeated with the electrostatic interactions turned off between the glycans. In the simulations of the vacuum condition, a stiffer harmonic spring with *k* = 5.0 kcal*/*(mol · Å^2^) was used for the biasing potential.

### Tetrahedral order parameter

We quantified the extent of tetrahedral geometry in the hydrogen bond network of water using the tetrahedral order parameter.^66^ For each water molecule, it is defined as,

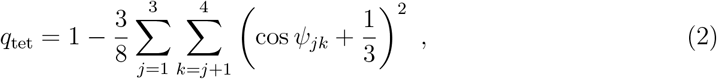

where *ψ*_*jk*_ is the angle between the lines joining the oxygen of the probe molecule and its *j*^th^ and *k*^th^ nearest heavy atoms which can donate or accept hydrogen bonds (that is, oxygen or nitrogen in water and glycan; see Fig. S7 for the illustration). For a perfect tetrahedral geometry, cos *ψ*_*jk*_ = −1*/*3 for all the angles between the nearest neighbors, resulting in *q*_tet_ = 1.

### Hydrogen bond dynamics

The time correlation function of a hydrogen bond (HB) is defined as,

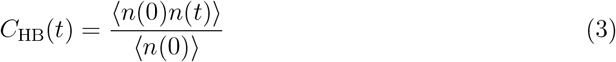

where *n*(*t*) is the hydrogen bond indicator for a given pair of HB-capable atoms at a given time step such that *n*(*t*) is 1 if hydrogen-bonded or 0 otherwise. The angular brackets denote an ensemble average over all the possible pairs (either water–water or water–glycan) throughout the simulation trajectory. Note that ⟨*n*^2^(*t*)⟩ = ⟨*n*(*t*)⟩ and thus the denominator in Eq. (3) normalizes the correlation function such that *C*_HB_(0) = 1. The lifetime or relaxation time of a hydrogen bond is calculated using 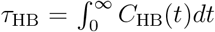. To define hydrogen bond formation, we adopted the criterion used elsewhere. ^105^

## Acknowledgment

We thank Guang Shi and Atreya Dey for useful discussions. We acknowledge the Texas Advanced Computing Center (TACC) at The University of Texas at Austin for providing computational resources that have contributed to the results reported within this paper. This work was supported by a grant from the National Science Foundation (CHE 2320256) and the Welch Foundation through the Collie-Welch Chair (F-0019).

## Supporting Information Available

The Supporting Information is available free of charge.

- Atomic partial charges on glycans (Fig. S1); orientation of ACE2 glycans (Fig. S2); comparison of hydrogen bond correlation functions (Fig. S3); hydrogen bond structure and kinetics of water near A4G4 (Fig. S4); effect of the solvent and electrostatics on the free energy profile for a glycan pair (Fig. S5); distributions of the distance between the first and the third sugar residues, and of the angle formed between two glycans (Fig. S6); illustration of the angle *ψ*_*ij*_ for the tetrahedral order parameter (Fig. S7)

## Notes

### Competing Interest Statement

The authors have declared no competing interest.

### Summary of Updates

Introduction and Discussion section were revised; Figure 4 added for the comparison to experiments.

