## Supplementary Information for "Water-mediated interactions between glycans are weakly repulsive and unexpectedly long-ranged"

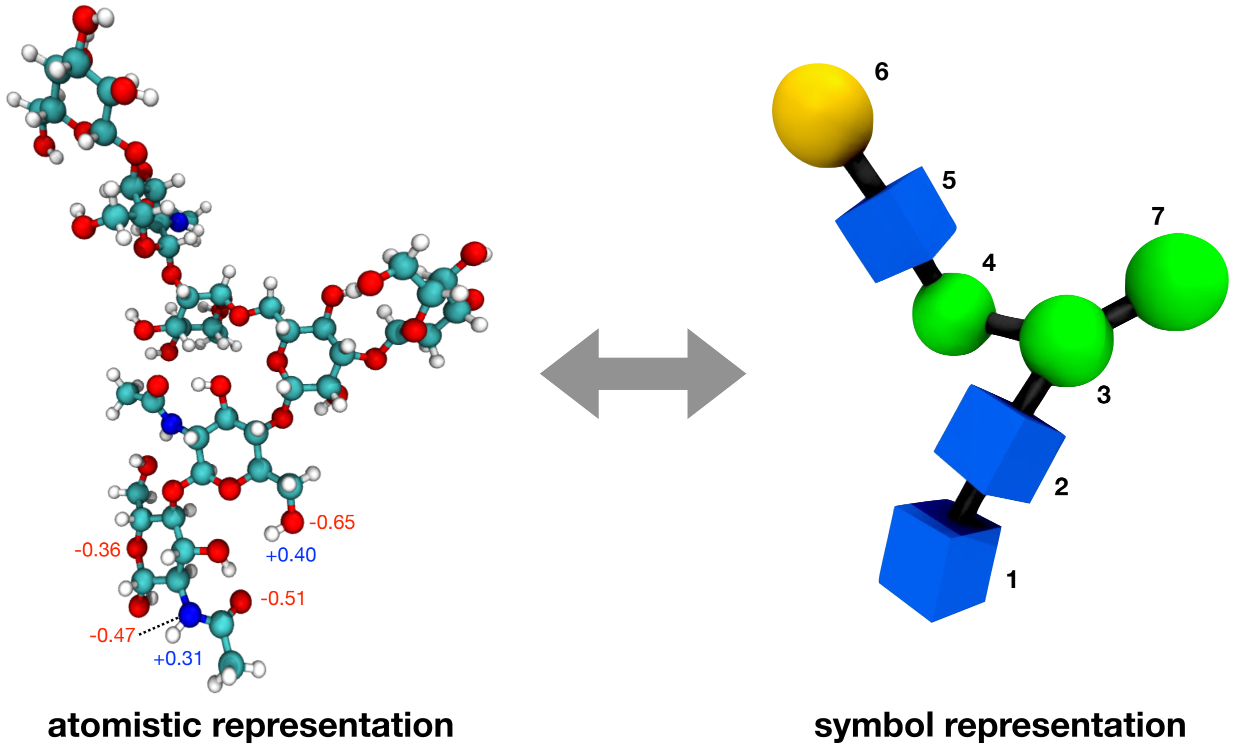


Figure S1: Glycans are capable of forming hydrogen bonds through polar atoms. (left) A1G1 glycan is shown in atomistic representation using balls and sticks, where carbon, nitrogen, oxygen, and hydrogen atoms are respectively in cyan, blue, red, and white colors. Different electronegative (oxygen, nitrogen) and electropositive hydrogen atoms are labeled with partial charges used in the molecular dynamics simulations based on the CHARMM36 force fields.[1] Note that TIP3P water,[2,3] the solvent used in the simulations, has the partial charges of −0.834 and +0.417 on oxygen and hydrogen atoms, respectively. (right) The glycan with the given structure is shown in a three-dimensional representation based on the Symbol Nomenclature For Glycans,[4,5] where the symbols representing the individual residues are located on the centers of the sugar rings. The labels are the residue numbers.

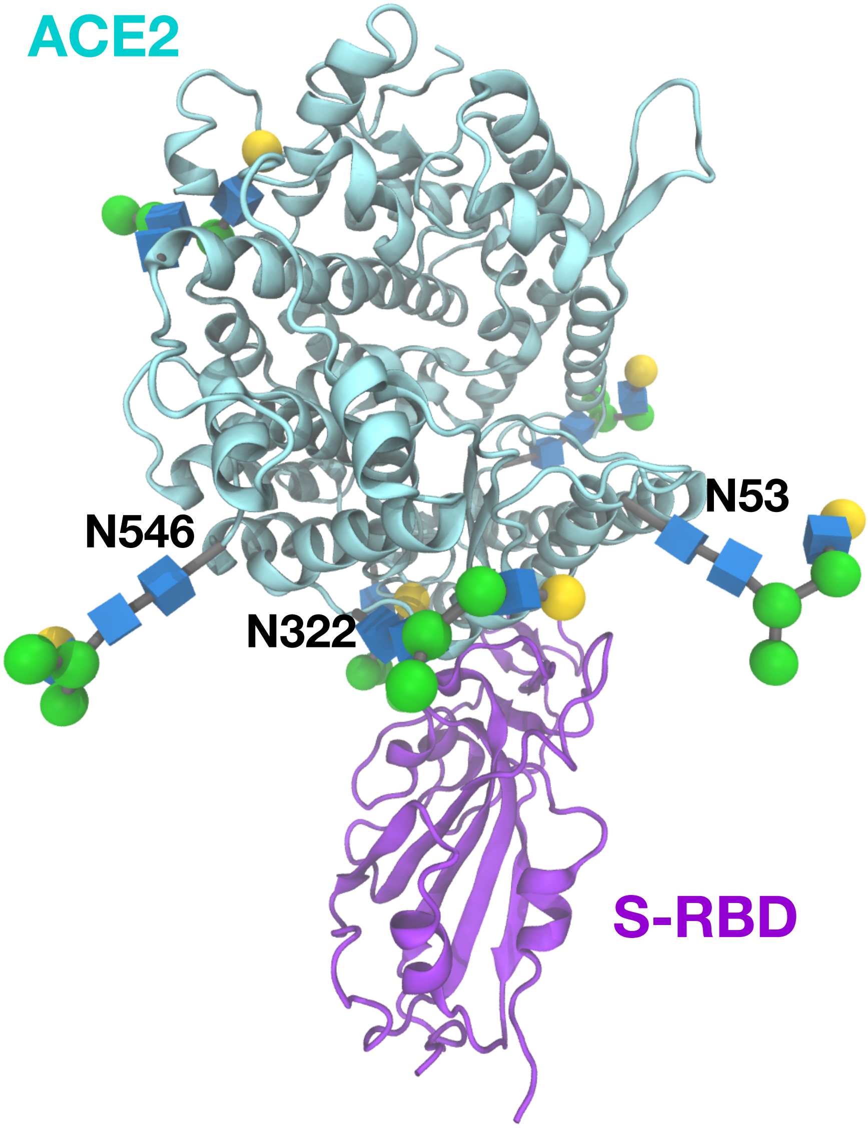


Figure S2: Illustration of glycan orientations in the glycosylated angiotensin converter enzyme 2 (ACE2) that is bound with the receptor binding domain (RBD) of the spike (S) protein of SARS-CoV-2. The glycans (A1G1 shown here) on three glycosylation sites (N546, N322, N53) in proximity are roughly parallel to one another. The structure of ACE2-RBD complex is based on the PDB: 6LZG.[6]

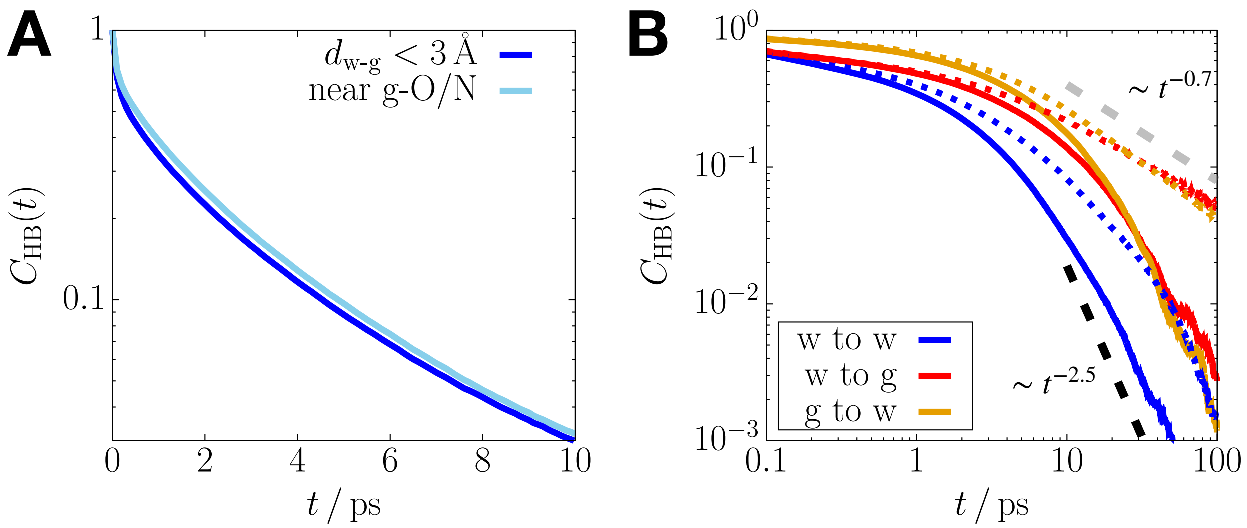


Figure S3: (A) Time correlation function of a hydrogen bond (HB) formed between water molecules in the hydration shell of a single glycan (blue), compared with those formed with water molecules near the oxygen or nitrogen atoms of the glycan (light blue). The $y$-axis is shown in a log scale. The difference between the correlation functions is not significant. (B) Log-log plot of the time correlation functions of a HB formed between water molecules (blue), donated from water to the glycan (red), and donated from the glycan to water (orange). The solid lines show the correlation functions for the HBs in the hydration shell of a single glycan, whereas the dotted lines are from the HBs of water in between the glycan pair constrained to $r = 7$ Å. The dashed lines show distinct power law relations for the long-time behavior of the correlation function.

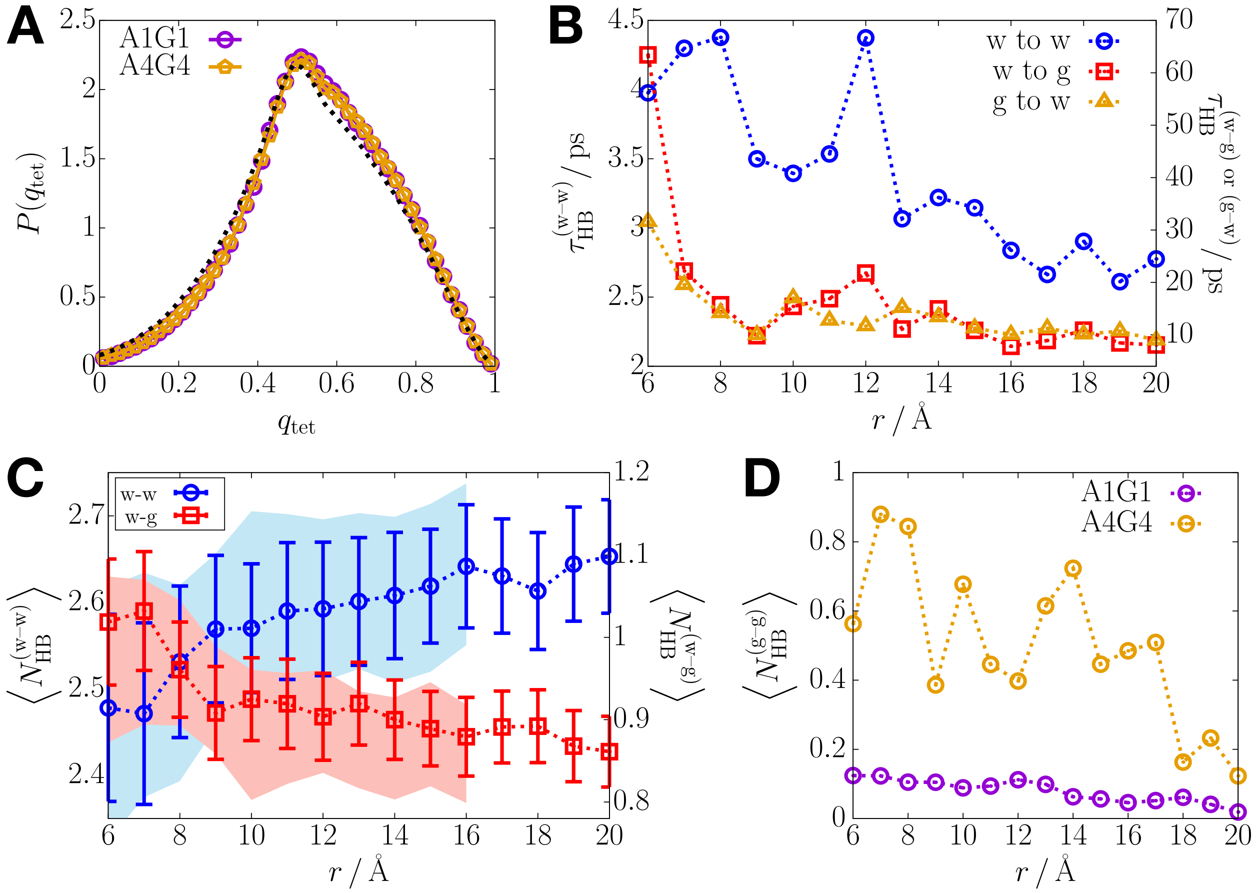


Figure S4: Hydrogen bond structure and kinetics for water molecules near A4G4. (A) Distribution of the tetrahedral order parameter for water's HB network in the hydration shell of a single A4G4 (orange), compared with that for a single A1G1 (purple). The black dotted line is for water molecules in between a pair of A4G4 constrained to $r = 13$ Å. (B) HB lifetime for water in between the A4G4 pair as a function of $r$. The left side of y axes corresponds to the values for the blue data points, whereas the right side is for the red and orange data. (C) Average number of HBs per water molecule in between the A4G4 pair as a function of $r$. The left and right sides of y axes correspond to the blue and red data points, respectively. For ease of comparison, the data for the A1G1 pair are shown in blue and red shades. (D) Average number of HBs formed between a pair of glycans as a function of $r$, shown in purple and orange for the A1G1 and A4G4 pairs, respectively.

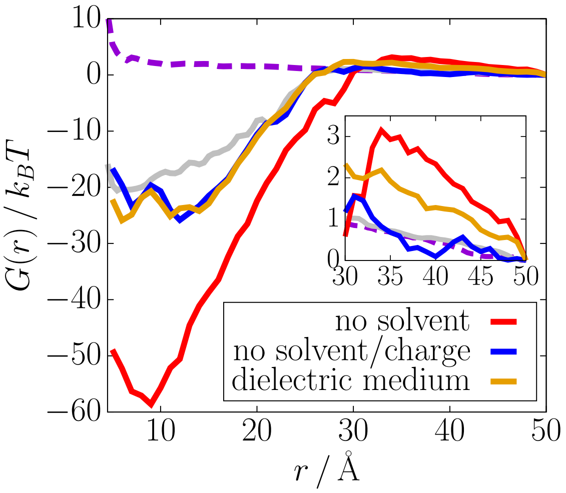


Figure S5: Effect of the solvent and electrostatic interactions on the free energy, $G(r)$, between the glycans. $G(r)$ for a pair of A1G1 in vacuum (red solid) is compared with the hydrated glycan pair (purple dashed). $G(r)$ for the uncharged glycans in vacuum and in the solvent are shown in blue and gray solid lines, respectively. The orange line corresponds to $G(r)$ for the fully charged glycans in a continuous dielectric medium with the relative permittivity of water (i.e., $\epsilon_r = 80$). These comparisons further underscore the importance of the electrostatic interactions associated with water whose dielectric constant is large.

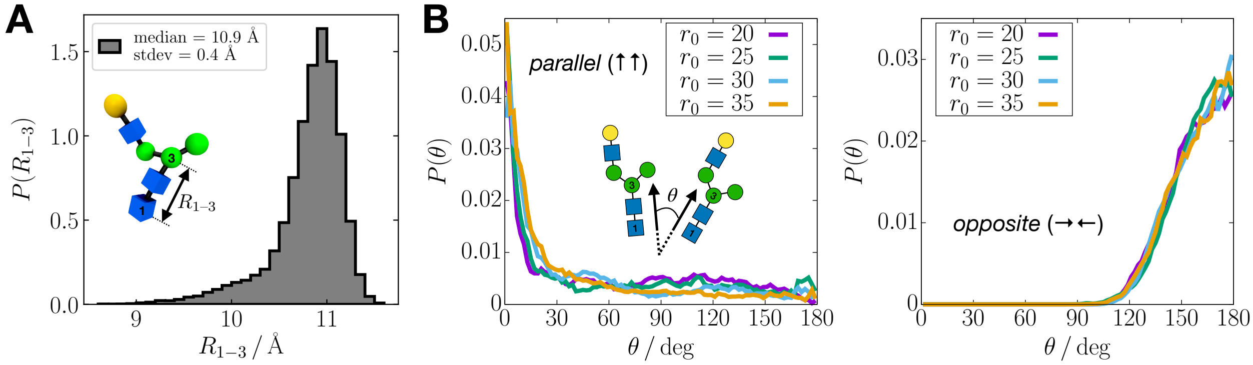


Figure S6: (A) Distribution of $R_{1\text{–}3}$, the distance between the centers of mass of the first and the third sugar residues of a single A1G1. (B) Distribution of the relative orientation between the A1G1 pair constrained in parallel (left) and opposite (right) directions (see Fig. 2A in the main text). Here the relative orientation is measured by $\theta$, the angle formed between the vectors, $\vec{R}_{1\text{–}3}$, from each glycan, as illustrated in the schematic.

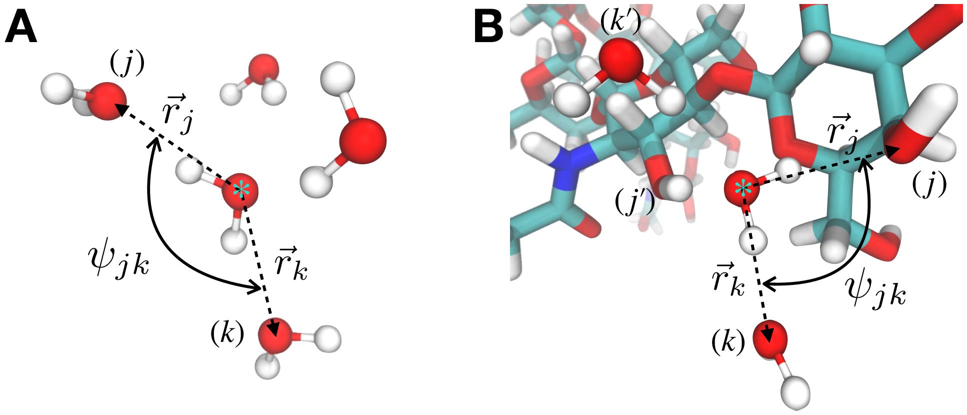


Figure S7: Illustration of the angle, $\psi_{jk}$, in Eq. (2) for calculating the tetrahedral order parameter, $q_{\text{tet}}$. (A) If a probe water molecule (marked with blue asterisk) is located far from a glycan, the four nearest water molecules that are closest to it are identified. Here $\vec{r}_j$ and $\vec{r}_k$ are the vectors from the oxygen atom of the probe water to the oxygen of the $j^{\text{th}}$ and $k^{\text{th}}$ neighboring water molecules, respectively. Then, the angle between two vectors is given by $\cos\psi_{jk} = (\vec{r}_j \cdot \vec{r}_k)/|\vec{r}_j||\vec{r}_k|$, where $|\vec{v}|$ is the length of the vector $\vec{v}$. (B) If a probe water molecule is adjacent to a glycan, the four nearest electronegative atoms, including the oxygen or nitrogen atoms in the glycan, are identified. The illustration shows the case in which the probe water (marked with blue asterisk) have hydrogen bonds with two hydroxyl groups in a glycan (marked as the $j^{\text{th}}$ and $j'^{\text{th}}$ neighbors) and two other water molecules (marked as the $k^{\text{th}}$ and $k'^{\text{th}}$ neighbors). From the positions of these electronegative atoms relative to the probe water's oxygen atom, $\psi_{jk}$ is calculated in the same way.
